# Acoustic features of emotional vocalisations account for early modulations of event-related brain potentials

**DOI:** 10.64898/2026.01.18.700181

**Authors:** Yichen Tang, Paul M. Corballis, Luke E. Hallum

**Author notes:** Contributing authors.

## Abstract

Emotion is key to human communication, inferring emotion in a speaker’s voice is a cross-cultural and cross-linguistic capability. Electroencephalography (EEG) studies of neural mechanisms supporting emotion perception have reported that early components of the event-related potential (ERP) are modulated by emotion. However, the nature of emotion’s effect, especially on the P200 component, is disputed. We hypothesised that early acoustic features of emotional utterances might account for ERP modulations previously attributed to emotion. We recorded multi-channel EEG from healthy participants (n = 30) tasked with recognising the emotion of utterances. We used fifty vocalisations in five emotions – anger, happiness, neutral, sadness and pleasure – drawn from the Montreal Affective Voices dataset. We statistically quantified instantaneous associations between ERP amplitudes, emotion categories, and acoustic features, specifically, intensity, pitch, first formant, and second formant. We found that shortly after utterance onset (120-250 ms, i.e., P200, early P300) ERP amplitude for sad vocalisations was less than for other emotional categories. Moreover, ERP amplitude at around 180 ms for happy vocalisation was less than for anger, sadness, and pleasure. Our analysis showed that acoustic intensity explains most of these early-latency effects. We also found that, at longer latency (220-500 ms; late P200, P300) ERP amplitude for neutral vocalisations was less than for other emotional categories. Furthermore, there were also ERP differences between anger and happiness, anger and pleasure, anger and sadness, happiness and pleasure, as well as happiness and sadness in shorter windows during this late period. Acoustic pitch and, to a lesser degree, acoustic intensity explain most of these later effects. We conclude that acoustic features can account for early ERP modulations evoked by emotional utterances. Because previous studies used a variety of stimuli, our result likely resolves previous disputes on emotion’s effect on P200.

## 1 Introduction

The human voice is an important and complicated element in interpersonal communication. Besides understanding the semantic content of speech, listeners infer a variety of information from the non-linguistic acoustic properties of voices, such as identity, age, gender, and the speaker’s current emotional state (Latinus & Belin, 2011). Human listeners can recognise the emotion in a speaker’s voice at rates significantly greater than chance, and this ability appears to be cross-cultural and cross-linguistic (Bachorowski, 1999; Scherer et al., 2001).

The neural mechanisms supporting recognition of emotion from emotional utterances remain unclear. Over the years, researchers have employed various neuroimaging methods to investigate this process (Beaucousin et al., 2007; Fecteau et al., 2007; Grandjean et al., 2006; Morris et al., 1999; Paulmann & Kotz, 2008; Pell et al., 2015; Pinheiro et al., 2017; Sander & Scheich, 2001; Schirmer & Kotz, 2006; Wildgruber et al., 2005). Electroencephalography (EEG) has been often used to study the temporal properties of this process due to its high temporal resolution. In studies examining emotional utterance perception using EEG, a series of event-related potentials (ERPs) time-locked to emotional utterance stimuli onsets have been observed. These include fronto-centrally distributed P50, N100, P200, P300, sometimes referred to as a “P200/P300 complex” (Schirmer & Gunter, 2017), and N400, as well as a centro-parietally distributed late positive potential (LPP), often occurring after 400 ms (Liu et al., 2012; Paulmann et al., 2013; Proverbio, De Benedetto, & Guazzone, 2020; Proverbio, Santoni, & Adorni, 2020; Toivonen & Rämä, 2009). Differences in ERPs between emotion categories or between emotional and neutral utterances have been detected as early as the P50 – i.e., 25-80 ms following utterance onset (Liu et al., 2012), and were also observed on N100 (Gädeke et al., 2013; Liu et al., 2012; Schirmer & Gunter, 2017), P200 (Gädeke et al., 2013; Liu et al., 2012; Paulmann & Kotz, 2008; Pell et al., 2015; Pinheiro et al., 2015; Schirmer & Gunter, 2017; Schirmer et al., 2013), P300 (Gädeke et al., 2013; Schirmer & Gunter, 2017; Wang et al., 2015), N400 (Proverbio, De Benedetto, & Guazzone, 2020; Proverbio, Santoni, & Adorni, 2020; Schirmer & Gunter, 2017), and LPP components (Schirmer & Gunter, 2017; Steber et al., 2020). However, the reported effects of emotion on these components, especially for the P200 component, are mixed. A single categorical factor like utterance type (e.g., sentences, words, and vocalisations) or participants’ attention to emotions does not easily explain the mixed results. For example, Pell et al. (2015) found a more positive-going P200 following happy than angry sentences, while Pinheiro et al. (2015) observed the opposite. Similarly, both Gädeke et al. (2013) and Steber et al. (2020) presented words uttered in different emotions while participants were not attending to the emotion. Gädeke et al. observed a more positive-going P200 following happy than neutral utterances, but Steber et al. again noted the opposite. Such controversies regarding the effects of emotion on early ERP components, especially P200, have been noted in several studies and reviews, yet no unified explanation has been established (Paulmann, 2023; Pell et al., 2015; Schirmer et al., 2013). Some of these studies have raised concerns that the amplitudes of these components are closely related to simple acoustic differences (Paulmann, 2023; Schirmer et al., 2013), however, none has conducted an in-depth analysis to address this issue. There are many reasons why the acoustic features of stimuli require special attention. Auditory evoked responses (AERs) share similar spatial and temporal distributions with early components of ERPs evoked by utterances, and these are known to be modulated by acoustic features such as intensity (Carrillo-de-la-Peña, 1999; Hegerl et al., 1994; Paiva et al., 2016; Sugg & Polich, 1995). Existing studies of emotional utterance perception have used stimuli of different type (e.g., vocalisations, words, sentences, etc.) and also custom-recorded stimulus sets, resulting in considerable variation in the acoustic properties of the stimuli (Gädeke et al., 2013; Liu et al., 2012; Paulmann & Kotz, 2008; Paulmann et al., 2013; Proverbio, De Benedetto, & Guazzone, 2020; Proverbio, Santoni, & Adorni, 2020; Schirmer & Gunter, 2017; Steber et al., 2020; Toivonen & Rämä, 2009). It would not be surprising if systematic acoustic differences between emotional stimuli could account for the observed effects attributed to emotion and explain the mixed results. In fact, it is generally accepted that differences in P50 and N100 reflect only systematic acoustic differences in emotional utterances (Paulmann, 2023; Schirmer & Kotz, 2006).

It is often believed that the acoustic features have been integrated in the first 200 ms during emotional utterance perception (Schirmer & Kotz, 2006). The subsequent ERP component, P200, reflects processing of emotionally “salient” features (Liu et al., 2012; Paulmann, 2023; Paulmann & Kotz, 2008; Pell et al., 2015; Schirmer & Kotz, 2006). N400 and LPP reflect cognitive evaluations of emotional properties such as valence (Paulmann et al., 2013; Proverbio, De Benedetto, & Guazzone, 2020; Schirmer & Gunter, 2017; Schirmer & Kotz, 2006; Steber et al., 2020). This ERP interpretation assumes that acoustic features are presented instantaneously at the utterance onset and processed once, overlooking the fact that an emotional utterance is a continuous, temporally unfolding sound. Recent studies make a different assumption and show that there are cortical responses time-locked to the temporally unfolding speech; these cortical responses continuously track acoustic features such as intensity and pitch (Bachmann et al., 2021; Ding & Simon, 2012; Lalor & Foxe, 2010; Teoh et al., 2019). Under this assumption, stimulus onset-locked ERPs reflect not only a discrete cortical response to an auditory onset, but also a convolution of time-locked cortical responses to acoustic features over preceding time windows (Lalor & Foxe, 2010).

While some studies have reported ERP differences between utterances sharing similar acoustic profiles, their computations of the acoustic profiles are frequently overly simplified. Most studies reported acoustic features averaged across entire stimulus recordings (e.g., average pitch and intensity of a stimulus on a given trial) instead of reporting the dynamics of acoustic features (e.g., changes of pitch or intensity across a given trial). Only Pell et al. (2015) reported pitch and intensity mean and standard deviation of the stimulus before key time windows. Since the acoustic features of emotional utterances may vary rapidly, summarising acoustic features across entire stimulus recordings leads to loss of temporal dynamics, and hence cannot accurately represent the acoustic properties the brain has processed before a given time window. Differences in a few key acoustic features may provide an alternative, direct explanation on the early emotion effects in ERPs.

We hypothesise that, the temporally dynamic acoustic properties of emotional utterances could account for apparent emotion effects in the first 500 ms of stimulus onset, encompassing P200 and P300. In particular, we expect acoustic intensity and pitch to be correlated with ERP amplitudes at a time lag; and we expect acoustic intensity and pitch to account for most of these emotional effects. Later time windows are excluded from the study due to potential bias introduced by systematic differences in stimulus duration between emotion categories (Section 2.2.2). In this study, we recorded EEG while presenting non-linguistic emotional vocalisations to participants. These vocalisations were drawn from a publicly available dataset (Belin et al., 2008). We chose these publicly available vocalisation utterances to exclude potential effects of linguistic prosody and semantic content on ERP components, as well as to ensure the reproducibility of our study. We analysed instantaneous associations between important acoustic features and ERP amplitudes following vocalisation onset. We also measured ERP differences and acoustic differences between emotions, and identified time windows where emotion’s putative effect on ERP amplitudes can be better explained by acoustics. We show that basic acoustic features, particularly intensity and pitch, could explain most emotion effects identified in our study and likely accounted for the mixed results observed in the existing literature.

## 2 Methods

### 2.1 Experiments

#### 2.1.1 Participants

Thirty participants (19 males and 11 females; aged 19 to 73, with an average age of 28.2±10.87 years, median 25.5 years) with normal hearing (self-reported) and normal or corrected-to-normal vision volunteered in our experiment. Each participant attended a single 3-hour session, before which the experimental procedure was explained, and written consent was obtained. Experimental protocols were approved by the University of Auckland Human Participants Ethics Committee (ref. UAHPEC24522). Subsequently, we removed four participants’ data due to a low signal-to-noise ratio (SNR), described in detail in Section 2.2.1. The two oldest participants (59 and 73 years) were included in the analysis. We also re-ran the analysis (Section 2.2.1) after excluding recordings from these two participants. The results were not different. Ultimately, we used data from 26 participants.

#### 2.1.2 Tasks and stimulus material

We tasked participants with listening to emotional vocalisations, and recognising the emotion of those vocalisations. Participants then reproduced the vocalisations naturally in the recognised emotion; we plan to present data pertaining to reproductions in a subsequent paper. We used the Montreal Affective Voices (MAV) dataset – a publicly available emotional vocalisation dataset —-as stimuli in this study (Belin et al., 2008). The MAV dataset has been experimentally validated to convey the labelled emotions. All vocalisations are based on the vowel /ah/ and contain no semantic information (Belin et al., 2008). We selected five emotion categories: happiness, anger, neutral, sadness, and pleasure, each consisting of ten vocalisations acted by five males and five females. A total of fifty emotional vocalisations were employed in this study. We chose the five emotions to cover both the high and low ends of the valence and arousal spectra. In particular, angry and happy vocalisations are both high in arousal but with negative and positive valences, respectively; pleasure and sadness are low in arousal but differ in valence; neutral vocalisations are low in arousal and neutral in valence (Belin et al., 2008).

#### 2.1.3 Experimental procedure

During the experiment, participants passively listened to emotional vocalisations, recognised the emotion category, and reproduced the /ah/ vowel in the recognised emotion. We explicitly asked the participants to focus on emotion and reproduce vocalisations in a natural and emotional way, while an exact mimic of vocalisations was not required (e.g. raising and dropping the pitch in the same way as the heard vocalisation). Each experimental session consisted of six blocks; each block consisted of 50 trials and 10 skippable practice trials. The fifty chosen vocalisations were randomly shuffled for each experimental block. Participant P18 attended only five blocks due to a lack of time.

Figure 1 illustrates the trial structure. Each trial started with participants fixating a central crosshair (width = 1° of visual angle) on a grey background for one second. Participants then listened to an emotional vocalisation followed by 0.5 s of silence. The participants recognised the emotion category of the vocalisation and then performed a reproduction task (these data are not reported in the present paper). Participants then chose the perceived emotion from a five-alternative forced-choice question, receiving feedback as well as the correct answer. Finally, the computer waited for a key press to proceed to the next trial. A running score displayed the number of completed and correctly answered trials in the top left corner of the computer monitor throughout each block. Participants were instructed to minimise movement and eye blinks, and maintain fixation. The experimental design for participants 15 through 30 was slightly different: for these participants, we fixed “T” (see Figure 1) to 3 s. Therefore, one of the fifty vocalisations that we used with a duration of 4.31 s was truncated. The mean vocalisation duration was 0.89 s. The average durations and standard deviations for the five emotion categories are reported in Table 1.

**Table 1:** The average durations and standard deviations for vocalisations in five emotion categories.

|  | Anger | Happiness | Neutral | Pleasure | Sadness |
| --- | --- | --- | --- | --- | --- |
| Duration (s) | $0.92 \pm 0.33$ | $1.45 \pm 0.51$ | $1.02 \pm 0.49$ | $1.35 \pm 0.44$ | $2.23 \pm 0.85$ |

**Fig. 1:**
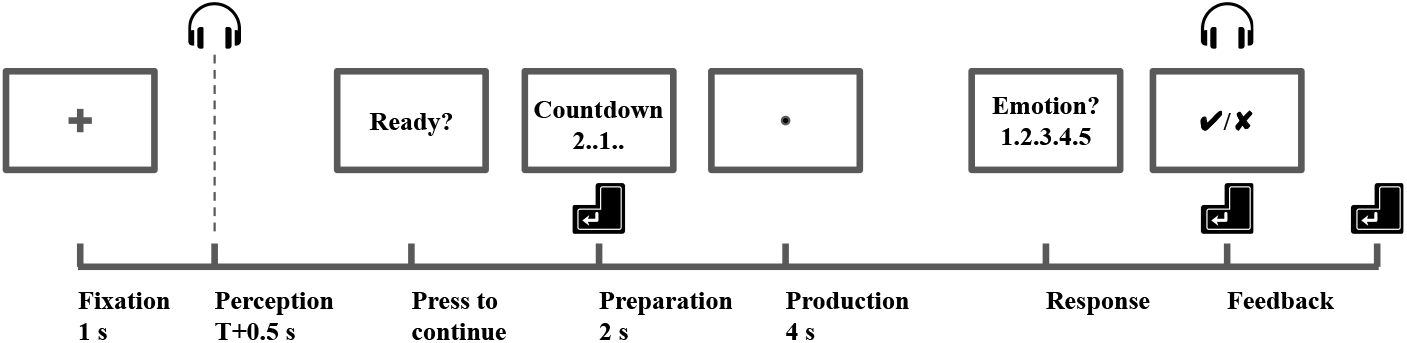
Illustration of a trial structure. The vocalisation stimuli were presented at the beginning of the Perception period. The duration of the Perception period depended on the duration of the stimuli, “T”. Prior to answering a five-alternative forced-choice question on the emotion of the vocalisation (“Response”), participants were tasked with reproducing the vocalisations. Here, we present analysis of vocalisation perception; we plan to present data pertaining to vocalisation reproductions in a subsequent paper.

#### 2.1.4 Visual and auditory setup

Visual stimuli were presented on a 24.5-inch OLED (organic light-emitting diode) display (PRM-224-3G-O; Plura Broadcast, Inc., Phoenix, Arizona, United States). Auditory stimuli, including vocalisations and auditory feedback (150ms pure tones at 800 Hz for correct answers and 200 Hz for incorrect answers) were delivered through a pair of insert earphones (30+ dB external noise exclusion; ER-1; Etymotic Research, Inc., Elk Grove Village, Illinois, United States). The earphones were calibrated before each experiment session to produce a 1 kHz tone at 70±1 dB when coupled to an occluded-ear simulator (IEC 60318-4:2010 Ed.1.0 standard; calibrated at 1 kHz to match a KEMAR simulator, G.R.A.S. Sound & Vibration, ApS, Holte, Denmark). We used a phototransistor (TEPT4400; Vishay Intertechnology, Inc., Malvern, Pennsylvania, United States) to record the onset of visual stimuli; we used a separate audio channel (iFi ZEN CAN amplifier, iFi Audio, Ltd., Southport, Sefton, United Kingdom) to record the onset of acoustic stimuli; these synchronisation signals were handled by Arduino UNO R3 boards.

#### 2.1.5 EEG and audio recordings

We recorded 64-channel EEG using a BioSemi ActiveTwo AD-box (ADC-17; ActiveTwo; Biosemi, B.V., Amsterdam, Noord-Holland, Netherlands) at a rate of 2048 samples per second. Scalp EEG electrodes were positioned according to the international 10-10 system using a BioSemi 64-channel headcap. The recorded channels included Fp1, AF7, AF3, F1, F3, F5, F7, FT7, FC5, FC3, FC1, C1, C3, C5, T7, TP7, CP5, CP3, CP1, P1, P3, P5, P7, P9, PO7, PO3, O1, Iz, Oz, POz, Pz, CPz, Fpz, Fp2, AF8, AF4, AFz, Fz, F2, F4, F6, F8, FT8, FC6, FC4, FC2, FCz, Cz, C2, C4, C6, T8, TP8, CP6, CP4, CP2, P2, P4, P6, P8, P10, PO8, PO4, and O2.

### 2.2 Data processing and statistical analysis

#### 2.1.1 EEG preprocessing

Offline preprocessing steps were applied using the open-source Python library MNE (Gramfort et al., 2013). Firstly, we applied a one-pass, zero-phase finite impulse response (FIR) filter to bandpass the EEG signals between 0.1 and 40 Hz. We interpolated bad channels, identified as shorted with the reference or ground electrodes, or containing excessive noise, using the spherical spline method (Perrin et al., 1989). We then re-referenced the channels to their average at each timestamp and removed electrooculography (EOG) contamination through the FastICA algorithm (Hyvarinen, 1999). We epoched the EEG signals from 0.2 s before to 4 s after the onset of the vocalisation stimuli, linearly detrended, resampled the data to 512 samples per second, and subtracted the baseline (0.2 to 0 s prior to stimuli onset). We excluded from further analysis those epochs which corresponded to trials wherein participants incorrectly recognised the vocalisation emotion. To improve the SNR, we clipped signals to the range between -40 and 40 µV. We then z-scored the signals across epochs, time, and channels, and outliers exceeding 3 standard deviations from the mean were again clipped. Finally, we averaged epochs corresponding to the same vocalisation stimuli, resulting in 50 averaged epochs for each participant. Participants P4, P8, P14, and P26 were excluded from the analysis due to poor SNRs, which resulted in non-visible P200 in their grand-averaged ERPs. This left a final sample of 26 participants included in the analysis.

#### 2.2.2 EEG and acoustic features

For each averaged epoch (Section 2.2.1), we extracted ERP amplitudes and acoustic features. ERP amplitudes were extracted by averaging amplitudes across every five timestamps (∼10 ms) from 0 to 0.5 s for each channel. The start-, central-, and end-times for each time window were recorded. We averaged channels within a chosen region of interest (ROI), specifically, across channels in the frontocentral region (AFz, F1, Fz, F2, FC1, FCz, FC2, C1, Cz, C2, CPz). This feature extraction process resulted in one value every 10 ms in each averaged epoch for each participant. We chose the ROI and the overall time window (0 to 0.5 s) to capture the amplitudes of early ERP components, including N100, P200, and P300 at 10 ms resolution. Later components, such as the LPP, were not analysed due to the overall short and unbalanced durations of vocalisations in the MAV dataset across emotion categories. Twelve, mostly angry or neutral, vocalisations lasted between 0.5 and 1 s, which would likely introduce bias to analyses on late ERPs. There were, however, only two vocalisations shorter than 0.5 s (one 0.24 s neutral and one 0.42 s angry vocalisation).

We surveyed the literature on EEG-based emotional utterance perception studies in healthy adults to establish a set of characteristic features for analysis. Among 33 studies mentioning acoustic features of the stimulus, duration, intensity, and pitch were most frequently noted, followed by formants mentioned in four studies (Charpentier et al., 2018; Paulmann et al., 2012; Schirmer et al., 2005; Wickens & Perry, 2015). Less frequent acoustic features included harmonic-to-noise ratio (HNR) (Charpentier et al., 2018; Paulmann et al., 2012) and jitter (Paulmann et al., 2012). In the broader literature, studies have also highlighted the importance of intensity, pitch, and HNR in human perception of emotional vocalisations (Anikin & Lima, 2018; Lima et al., 2013). Our analysis (see Section 2.2.3) examines the effects of each individual acoustic feature separately to maintain simplicity and comprehensiveness. Therefore, we restricted ourselves to examine only the most noted features – duration, intensity, pitch, and formants. The duration is not a temporal acoustic feature and is therefore excluded from this study. Among all formants, we selected the first two, as in Wickens and Perry (2015). We also examined the HNR. However, statistically significant differences in instantaneous HNR were observed only between a few emotion pairs and were mostly confined to short time intervals (see Appendix B). Therefore, HNR was excluded from subsequent analyses.

We extracted four acoustic features from the corresponding vocalisation stimuli: intensity, pitch (i.e., F0, fundamental frequency), first and second formants (F1 and F2). These features were computed with a step size of 10 ms, matching the time windows for ERP measurements. We used Praat (Boersma & Weenink, 2021) interfaced in Python through the open-source library Parselmouth (Jadoul et al., 2018) to extract these acoustic features. For pitch extraction, we set parameters in Praat as follows: 75 Hz pitch floor, 600 Hz pitch ceiling, 0.25 voicing threshold, 0.03 silence threshold, 0.55 octave jump cost, and 0.28 voiced-unvoiced cost (Boersma & Weenink, 2021). The pitch estimation parameters were carefully tuned, and errors in individual pitch contours were manually corrected using Praat’s built-in “View & Edit” function, following guidelines by Styler (2013) and Praat (Boersma & Weenink, 2021). The extracted and corrected pitch contours for all 50 vocalisations are plotted in Appendix A.

#### 2.2.3 Statistical analyses for identifying acoustically-explainable ERP differences between emotion pairs

We hypothesised that there are points in time where the underlying systematic acoustic differences drive the ERP differences between emotion categories and explain the putative emotion effects on ERP amplitudes, as illustrated in Figure 2. In these cases, the acoustic features serve as the most parsimonious explanatory factor accounting for both the categorisation of emotion and the corresponding ERP modulations. We verified this hypothesis using statistical tests in 10 ms windows between 0 and 500 ms following stimulus onset. We performed the following statistical tests examining the associations among acoustic features, emotion categories and ERP amplitudes:

**Fig. 2:**
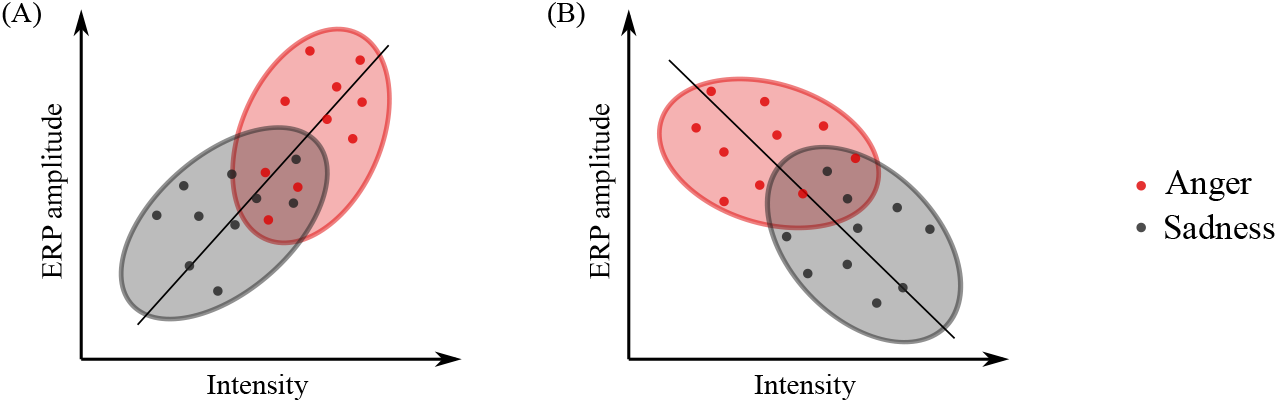
Systematic differences in vocalisations’ acoustic features between emotion categories may drive ERP differences, which appear to be an emotion effect. For example, subplot (A) illustrates a sample situation where at a timestamp, acoustic intensity is positively correlated to the ERP amplitude. Furthermore, angry vocalisations have a higher acoustics intensity than sad vocalisations. The ERP differences between angry and sad vocalisations may appear to be an emotion effect, however, in this case, it could be explained by acoustic intensity. Similarly, subplot (B) illustrates a different situation where a negative correlation between acoustic intensity and ERP amplitude could also explain ERP differences between anger and sadness.

1. We computed the instantaneous correlation between each acoustic feature – intensity, pitch, F1, F2 – and ERP amplitude (“Acoustics ∼ ERP”, here, ∼ denotes an association between two factors).
2. For each acoustic feature, we computed instantaneous pairwise differences between emotion categories (“Acoustics ∼ Emotion”).
3. We computed differences in ERPs between emotion categories (“Emotion ∼ ERP”).

We assumed that the cortical response to the acoustic features comprising these short vocalisations was consistently lagged as the vocalisations unfold, following recent studies that have found timelocked cortical responses to acoustic features unfolding over time (Bachmann et al., 2021; Ding & Simon, 2012; Lalor & Foxe, 2010; Teoh et al., 2019). We found that a lag value of 100 ms maximised the correlation between acoustic features and ERP amplitudes and hence we adopted 100 ms as the lag in the following analyses. This 100 ms lag is also consistent with the latency of the estimated cortical response peak reported by Lalor and Foxe (2010) and Ding and Simon (2012).

We conclude that an acoustic feature could explain an ERP difference between two emotion categories at a timestamp if all three statistics at the timestamp reached statistical significance and the associations followed the two patterns illustrated in Figure 2. This process can be simplified to a sequence of yes/no questions:

1. For ERP amplitude differences (eg., happiness versus anger) at time *t*, are there differences in acoustic features (eg., happiness pitch versus anger pitch) at *t* − 100 ms?
2. If so, is there a correlation between the corresponding acoustic feature (eg., pitch) at *t* − 100 ms and the ERP amplitude at *t*, regardless of emotion?
3. If so, is the sign of the correlation consistent with both the ERP amplitude differences and the acoustic feature differences?

As an example, if we observed a more positive-going ERP for anger than happiness, a higher pitch for anger than happiness, as well as a positive (but not negative) correlation between pitch and ERP amplitude, then we would conclude that pitch accounts for this ERP difference between anger and happiness. In the following paragraphs, we detail the three statistical tests mentioned above, as briefly illustrated in Figure 3.

**Fig. 3:**
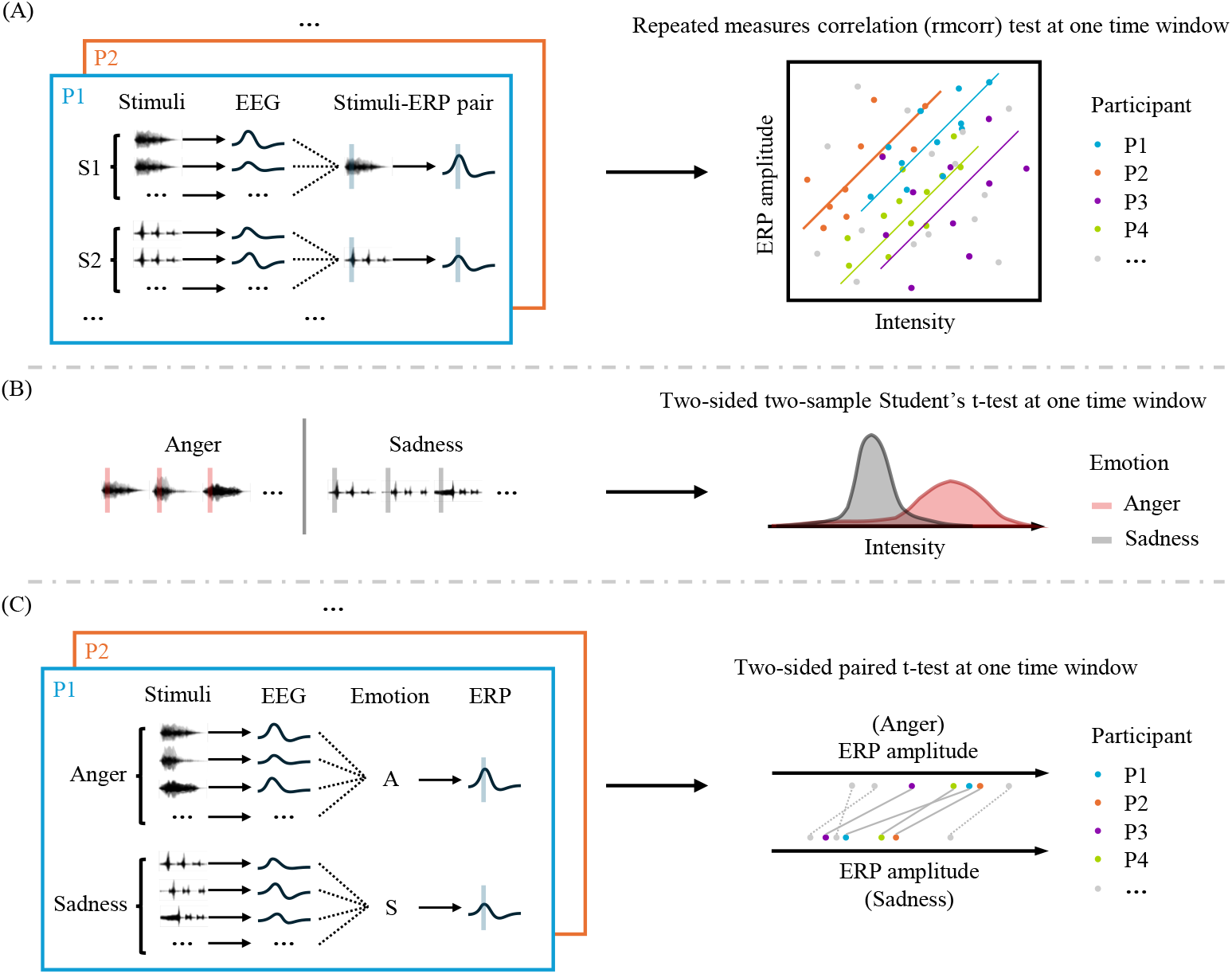
We computed three statistics at each 10 ms window, examining the associations among the acoustic features, emotion categories and ERP amplitudes. Subplot (A) illustrates a test performed to examine the correlation between acoustic intensity and ERP amplitude (“Acoustics ∼ ERP”). EEG epochs for the same stimulus and participant were averaged to form stimuli-ERP pairs. Acoustic intensity and ERP amplitudes were extracted from the target time window and a repeated-measures correlation (rmcorr) test was performed to examine the correlation across participants. Subplot (B) illustrates a test performed to examine acoustic intensity differences between angry and sad vocalisations (“Acoustics ∼ Emotion”). A two-sided two-sample Student’s t-test was performed on the acoustic intensity extracted from all angry and sad vocalisations over the target time window. Subplot (C) illustrates a test performed to examine ERP differences following angry and sad vocalisations (“Emotion ∼ ERP”). EEG epochs for stimuli of the same emotion and the same participant were averaged to form emotion-ERP pairs. ERP amplitudes were extracted from the target time window and a two-sided paired t-test was performed to examine the emotion effect across participants. For all statistical tests, we employed a cluster-wise non-parametric permutation test to control for false positives (Maris & Oostenveld, 2007).

##### Acoustics ∼ ERP

We computed correlations between acoustic feature measurements and ERP amplitudes (Figure 3-A). We averaged EEG signals in each participant across sessions to obtain 50 ERP responses, one for each vocalisation stimulus. We assumed that the brain would produce a consistent response to the same vocalisation across sessions, such that averaging the responses reduces unwanted variability and noise. Next, for each acoustic feature, we obtained its measurements for the 50 vocalisations respectively. Then, we performed repeated measures correlation (rmcorr) tests between this acoustic feature and the ERP amplitude, separately for each 10ms window (Bakdash & Marusich, 2017). For example, from each ERP response, we computed its window averaged value from 100 to 110 ms following stimulus onset; and from each vocalisation stimulus, we computed the intensity measurement from 0 to 10 ms, respectively. We then applied the rmcorr test on this sample of 50×26 participants = 1300 intensity-ERP measurement pairs, treating the 50 ERPs and intensity measurements as the repeated measures for each participant. We repeated the above process on all four characteristic acoustic features (i.e. intensity, pitch, F1 and F2) and for all time windows between 100 and 500 ms.

To control for false-positives, we employed a cluster-wise non-parametric permutation test (Maris & Oostenveld, 2007). We first combined statistically significant *r* values (p≤0.05) that were temporally adjacent and have the same sign into clusters. We then computed cluster-wise statistics by summing up the statistics within each cluster. We obtained a null distribution of cluster statistics by repeating the process 1000 times after randomly shuffling ERP measurements across epochs and participants. Finally, we computed a p-value for each observed cluster by comparing the observed cluster statistics with the null distribution. Since there were four acoustic features, we used a critical value of 0.05*/*(4 features) = 0.0125 to determine if the observed p-values were statistically significant. To further confirm ERP-Acoustic correlations, we also tested acoustic features’ correlations with ERP amplitudes within each emotion category. For this analysis, we performed the cluster-wise non-parametric permutation test on all test results for five emotion categories and applied a critical value of 0.05*/*(4 features × 5 emotions) = 0.0025.

##### Acoustics ∼ Emotion

We computed differences in vocalisation acoustic features between emotion pairs (Figure 3-B). We first targeted one acoustic feature and retrieved its measurements on the 50 vocalisation stimuli. Then, we performed two-sided two-sample Student’s t-tests on the measurements between each pair of emotions in 10 ms windows. For example, we computed the mean intensity in the first 10 ms for each vocalisation stimuli. Then, we performed the t-test between ten angry and ten sad vocalisations’ intensity measurements. We note that the statistics here for the 0 to 10 ms window were analysed together with the other two statistics for the time window between 100 and 110 ms. We repeated the process for all emotion pairs, characteristic acoustic features, and time windows between 0 and 400 ms. We applied the same cluster-wise non-parametric permutation test, with a critical value of 0.05*/*(4 features×10 emotion-pairs) = 0.00125. We performed pairwise t-tests between emotions such that the direction of the differences can be derived for further comparison with the other statistics. The direction of the differences was also considered in the cluster-wise non-parametric permutation test when clustering the statistics.

##### Emotion ∼ ERP

We computed differences in ERP amplitudes between emotion pairs (Figure 3-C). We first averaged EEG signals across epochs within the same emotion category for each participant, resulting in five ERP responses, one for each emotion. This corresponded to a total of 5×26 participants = 130 ERP responses. We then performed two-sided paired t-tests in each 10 ms time window. For example, we computed the mean amplitude between 100 and 110 ms for each ERP response and performed the paired t-test between anger and sadness (i.e. 26 anger ERP amplitude measurements v.s. 26 sadness ERP amplitude measurements, one pair of measurements per participant). We repeated the process for all emotion pairs and for all time windows between 100 and 500 ms. Again, we applied the cluster-wise non-parametric permutation test, using a critical value of 0.05*/*(10 emotion-pairs) = 0.005.

## 3 Results

### 3.1 Behaviour results

Behavioural analysis showed that participants capably performed the emotion recognition task and achieved low error rates, as tabulated in Table 2. Overall, participants incorrectly recognised emotion in a total of 293 (3.76%) of 7800 trials. Participants made more errors when attempting to recognise angry and pleasant vocalisations than happy, neutral, and sad vocalisations. Participants also tended to confuse angry vocalisations with pleasant vocalisations, and confuse pleasant vocalisations with angry and neutral vocalisations, as shown in Figure 4. The error trials were removed from further analyses.

**Table 2:**
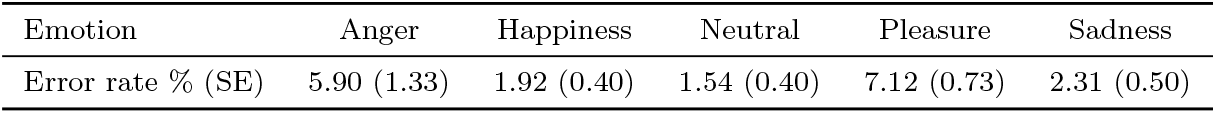
Behavioural error rates (%) averaged across participants and their standard errors (SE), for the task of recognising the emotion of vocalisations.

| Emotion | Anger | Happiness | Neutral | Pleasure | Sadness |
| --- | --- | --- | --- | --- | --- |
| Error rate % (SE) | 5.90 (1.33) | 1.92 (0.40) | 1.54 (0.40) | 7.12 (0.73) | 2.31 (0.50) |

**Fig. 4:**
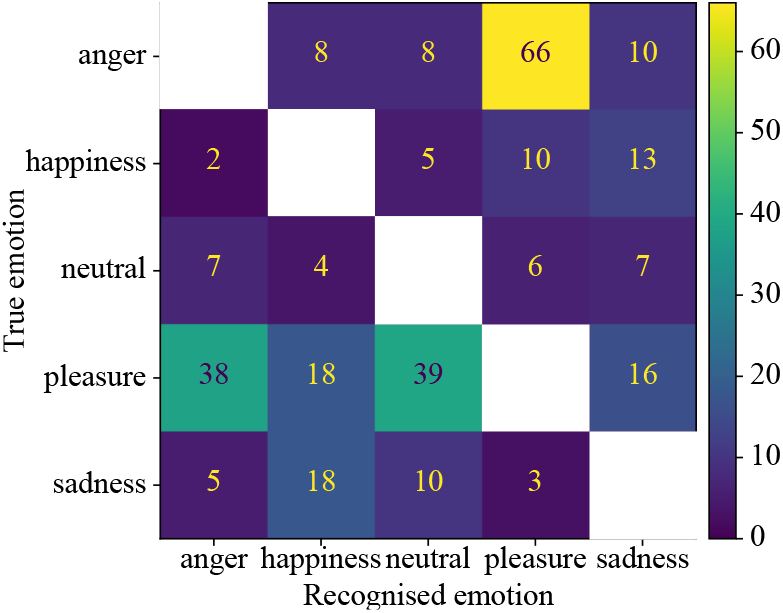
Confusion matrix showing performance pooled across all participants tasked with recognising emotions of vocalisations. Participants tended to confuse angry with pleasant vocalisations, and pleasant with angry or neutral vocalisations. The numbers represent the total number of vocalisations in one emotion that were confused with another. For example, a total of 66 angry vocalisations were incorrectly recognised as pleasant. The number of correctly recognised trials (i.e., the matrix diagonal) is omitted.

### 3.2 ERPs and acoustic features

We computed the average ERP response to vocalisation in each of our five emotion categories; here, we pooled across participants and channels comprising our ROI. These ERP responses showed clear N100, P200 and P300 components, as shown in Figure 5-A. The N100 peaked between 0.1 and 0.13 s following stimuli onset (0.110 s for anger, 0.106 s for happiness, 0.101 s for neutral, 0.110 s for pleasure, and 0.126 s for sadness); the P200 peaked between 0.2 and 0.24 s (0.214 s for anger, 0.231 s for happiness, 0.202 s for neutral, 0.204 s for pleasure, and 0.235 s for sadness); the P300 peaked between 0.3 and 0.36 s and was most pronounced for happiness (0.358 s), neutral (0.329 s), and sadness (0.329 s). We also computed the characteristic acoustic features (i.e., intensity, pitch, F1, and F2) in 10 ms windows. The acoustic features showed rich temporal dynamics, as exemplified in Figure 5-B. Notably, these rich temporal dynamics would have been lost had we simply averaged across the vocalisation period.

**Fig. 5:**
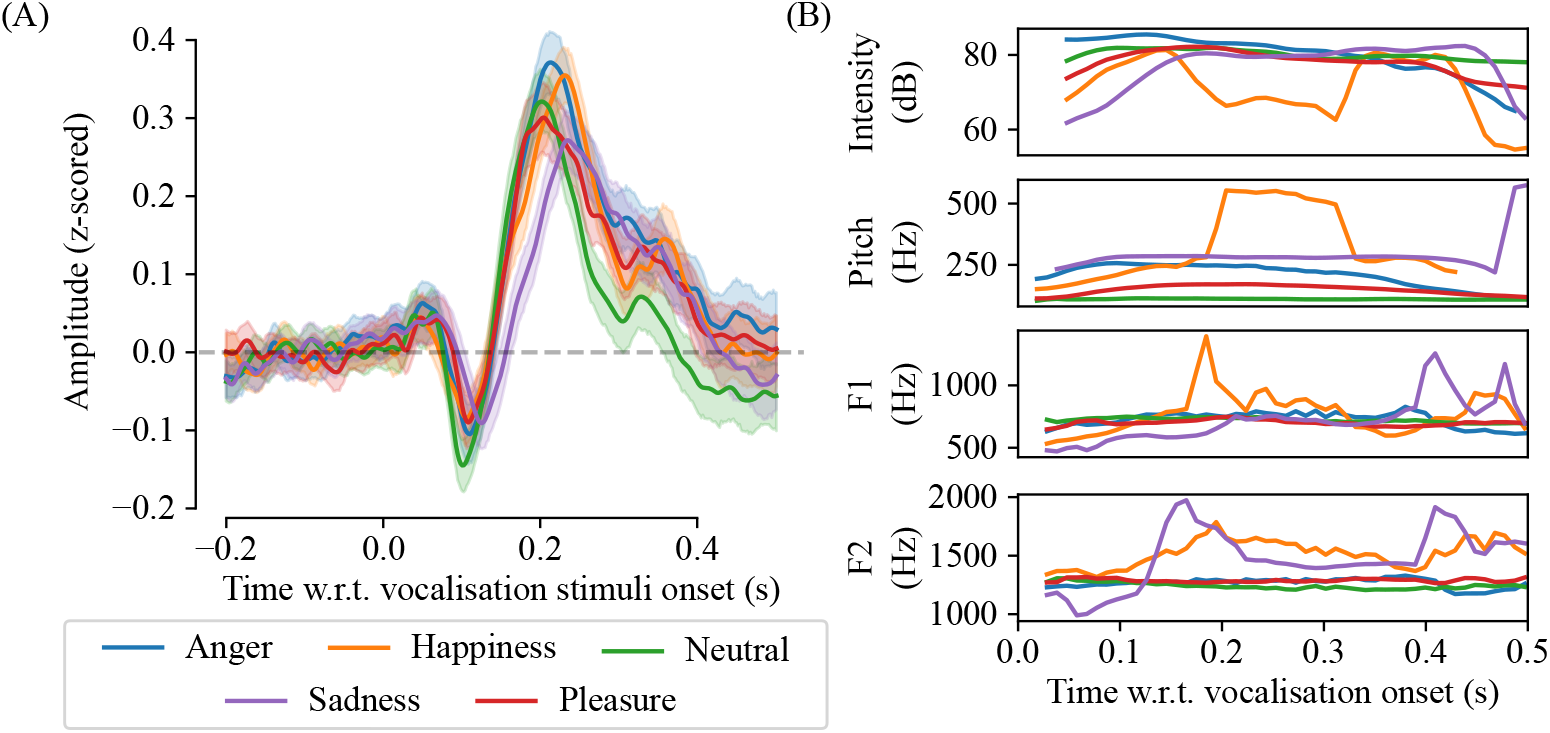
Event-related potentials (ERPs) were prominent in EEG recordings following vocalisation onset. Both ERPs and acoustic features show rich temporal dynamics that differ with emotion category. The plot in (A) shows the averaged ERP from -0.2 to 0.5 s following vocalisation onset. These ERP responses represent an average across 11 fronto-central channels in our region of interest (i.e., AFz, F1, Fz, F2, FC1, FCz, FC2, C1, Cz, C2 and CPz) and 26 participants. The shaded areas represented the 95% confidence intervals of the ERP amplitudes across participants. The plots in (B) show sample acoustic features computed from actor number 55 in the MAV dataset. Intensity, pitch, first formant (F1), and second formant (F2) were computed using Praat (Boersma & Weenink, 2021) for each vocalisation stimuli every 10 ms.

### 3.3 Statistical analysis results

#### 3.3.1 Instantaneous correlations between acoustic features and ERP amplitudes

We first examined the correlations between vocalisation acoustics and the ERP amplitudes in 10 ms windows. Here, we assumed a fixed lag from when the stimuli reached the participants’ ears to when the ERPs were reflected on the participants’ scalps. We found that a lag value of 100 ms maximised the Acoustics-ERP correlations and hence we adopted 100 ms as the lag in the following analyses.

The four acoustic features were statistically significantly correlated with the ERP amplitudes across a total of 99 time windows (∼24 windows per feature) from 140 to 500 ms after vocalisation onset, as shown in Figure 6-A. Correlations were also observed within emotion categories, as shown in Figure 6-B. Notably, during 150 to 210 ms, positive correlations with ERP amplitudes were observed for all acoustic features, both across emotion categories and within emotions. Particularly, there was a relatively high positive correlation between intensity and ERP amplitudes during 140 and 220 ms 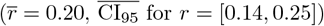, which was also reflected specifically within neutral and pleasure emotion categories. When data were pooled across emotion categories, pitch was positively correlated to the ERP amplitudes at most points in time (Figure 6-B; 150 to 330 ms, 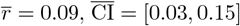; 340 to 500 ms, 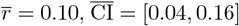), despite its showing relatively low correlation for individual emotions (Figure 6-B).

**Fig. 6:**
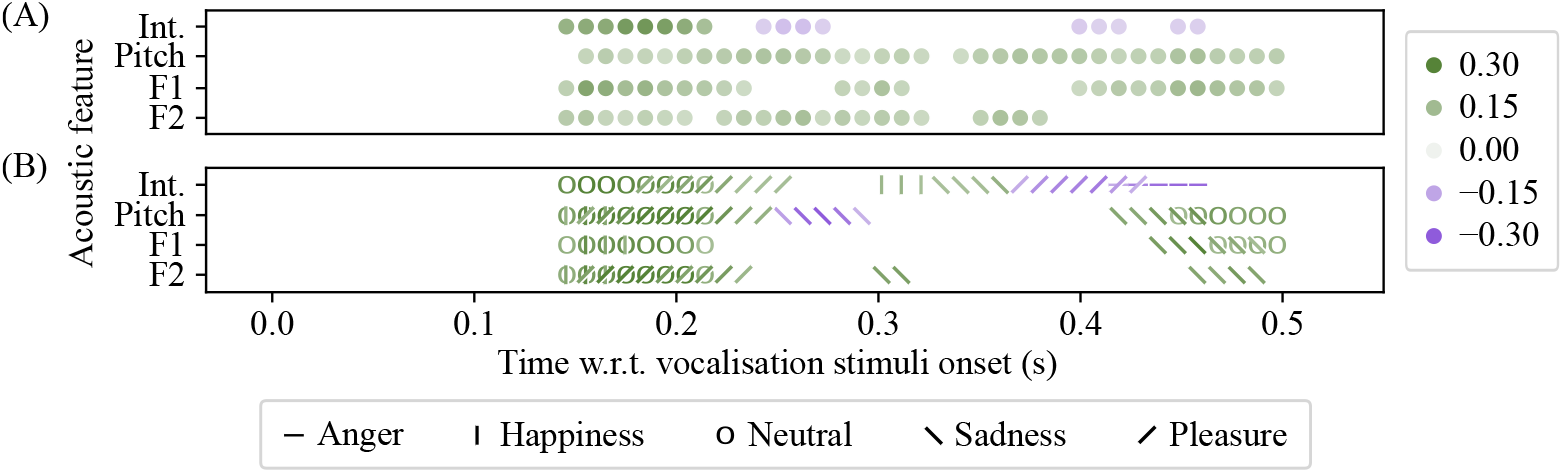
There were instantaneous correlations between acoustic features and ERPs. Plot (A) shows the points in time where the ERPs were statistically significantly correlated with each acoustic feature (intensity, pitch, F1 and F2; lag = 100 ms). All acoustic features were correlated with the ERP amplitudes. For example, the intensity was positively correlated to ERP amplitudes during 140 to 220 ms (averaged *r* statistics, 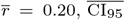 for *r* = [0.14, 0.25]; greater intensity predicts more positive-going ERP amplitudes). Plot (B) shows the points in time where the ERPs were statistically significantly correlated with each acoustic feature, separately for each emotion (distinguished by different marker shapes). For example, F1 was positively correlated to ERP amplitudes during 140 to 220 ms 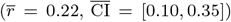 for emotions happiness and neutral, and during 440 to 500 ms 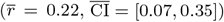 for emotions sadness and neutral.

#### 3.3.2 Instantaneous differences in acoustic features between emotion category pairs

We then computed differences in acoustic features between emotion pairs. There were clear differences in intensity and pitch between pairs of emotions, as plotted in Figure 7. Intensity showed the greatest difference (186 time windows over 9 emotion pairs) between emotion, and the highest t-statistics 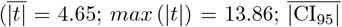 for mean intensity difference = [5.19, 15.66] dB). Notably, sad and happy vocalisations tended to have a lower acoustic intensity than angry, pleasant, and neutral vocalisations between 40 and 350 ms. Pitch was statistically significantly different in 161 time windows across eight emotion pairs 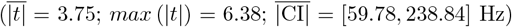. Notably, neutral vocalisations tend to have a lower pitch than all other vocalisations; angry vocalisations also tend to have a higher pitch than others. The F1 and F2 features were less different between emotions (F1, 57 windows, 6 pairs, 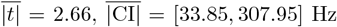 F2, 84 windows, 5 pairs, 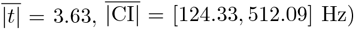, and were found less important for explaining ERP differences which is discussed later in Section 3.3.3, hence, details for F1 and F2 differences between emotions are not presented here.

**Fig. 7:**
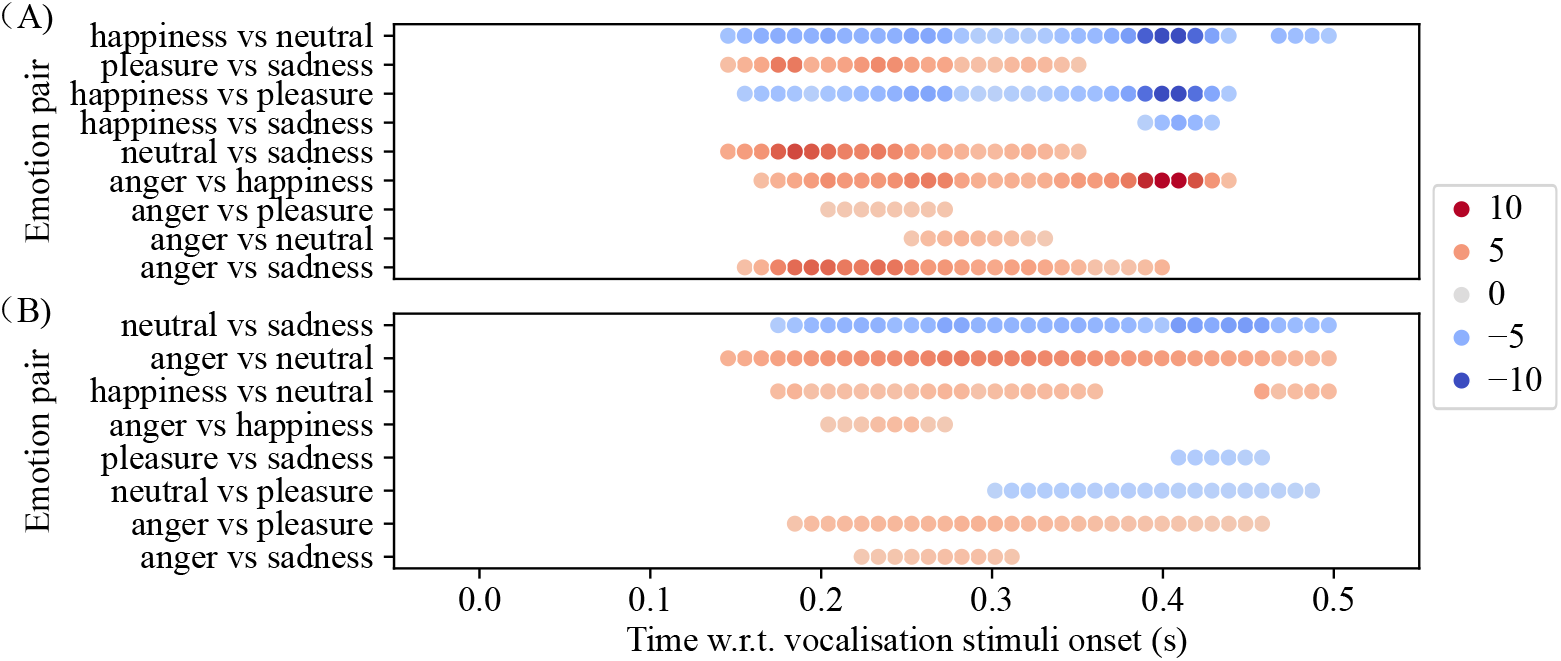
There were instantaneous differences in acoustic features between emotional category pairs. Plot (A) shows, between pairs of emotions, the points in time where there were statistically significant differences in acoustic intensity. The red/blue colour map represents the value of the *t* statistic (see Methods). Here, we adjusted the x-axis to account for the 100 ms lag used. For example, the intensity of pleasant vocalisations was statistically significantly greater than that of sad vocalisations during 140 to 360 ms 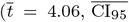 for mean intensity difference = [3.99, 14.53] dB). Emotion pairs with no statistically significant differences in acoustic features are omitted. Plot (B) shows, between pairs of emotions, the points in time where there were differences in acoustic pitch. For example, the pitch of neutral vocalisations was less than that of the sad vocalisations during 170 to 500 ms 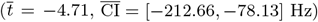.

#### 3.3.3 Instantaneous differences in ERPs between emotion pairs can mostly be explained by acoustics

Finally, we computed ERP differences between emotion pairs and determined where the differences were statistically significant. We then identified points in time where these ERP differences could be explained by acoustic features. For example, shown in Figure 8, angry vocalisations evoked more positive-going ERPs than happy vocalisations at 190 ms; we then found that intensity for angry vocalisations was also greater than intensity for happy ones, similarly we found that intensity was positively correlated to the ERP amplitudes, at the corresponding time point (190 ms). Thus, acoustic intensity could explain the ERP differences between anger and happiness at 190 ms. Note that in the above example, if the directions of the differences and correlations did not match (e.g., there was a negative intensity-ERP correlation), then we do not claim explainability. A total of 163 time windows across ten emotion pairs were found with statistically significantly different ERP amplitudes between emotions, while most (132 out of 163, 81.0%) of these time windows were explainable by one or more of the acoustic features (intensity, pitch, F1 and F2). Specifically, 121 (74.2%) of these time windows were explainable by intensity and pitch. Figure 8 plots the statistically significant ERP differences between emotion pairs; here, we use symbols (+,×) to represent time points at which intensity and pitch can explain ERP differences, and we use circles to represent time points at which no acoustic feature can explain ERP differences.

**Fig. 8:**
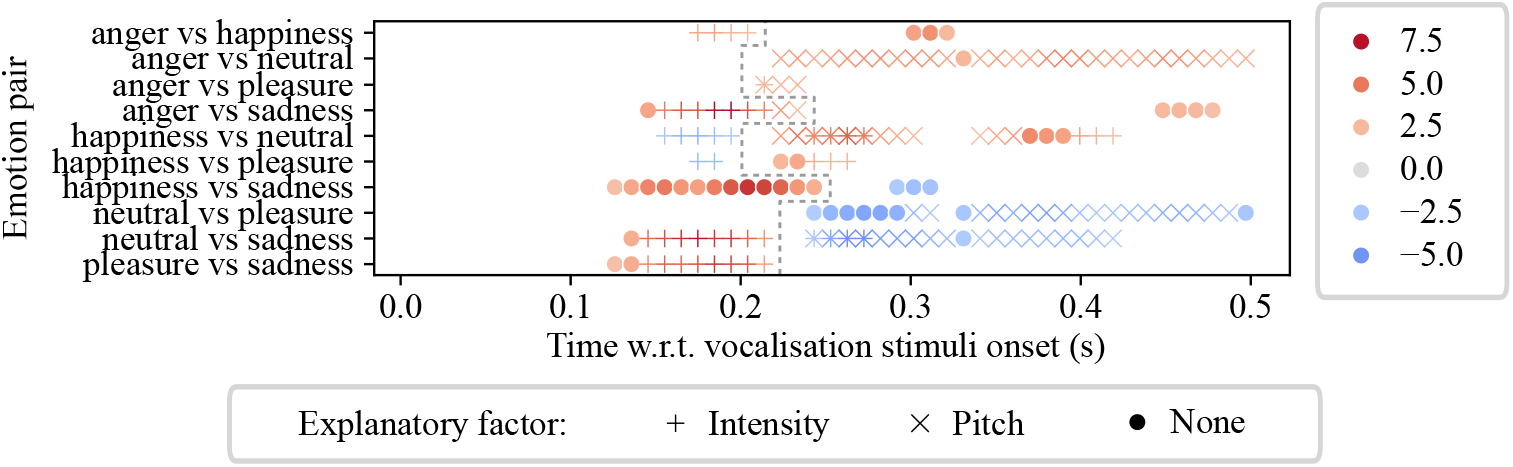
Instantaneous differences in ERPs between emotional category pairs are, at times, explained by acoustic features intensity and pitch. The plot shows, for several pairs of emotions, the points in time where there was a statistically significant difference between ERPs. The symbols mark where this difference can be explained by intensity (+), pitch (×); the circles mark points in time where neither of these acoustics features can explain the difference. For example, across all speakers, angry vocalisations evoked more positive-going ERPs during intervals 170 to 210 ms and 300 to 330 ms as compared to happy vocalisations. The differences between 170 and 210 ms could be explained by systematic differences in the intensity of vocalisations, and the pitch of vocalisations (at 200 ms only). The differences during interval 300 to 330 ms, however, could not be explained by either intensity or pitch. We use the dashed line to delineate early (clusters starting before 200ms) and late (clusters starting after 200ms) times of interest. In early clusters, intensity tended to explain most ERP differences, while in late clusters, pitch tended to explain more ERP differences.

The analysis we show in Figure 8 revealed marked differences between the effect of intensity and that of pitch: early ERP differences (120 to 250 ms, that is, clusters starting before 200 ms) were mostly explained by intensity, while late ERP differences (220 to 500 ms, that is, clusters starting after 200 ms) were mostly explained by pitch, and only partially by intensity. In early clusters, sad vocalisations evoked less positive-going ERPs than other vocalisations between 150 and 220 ms 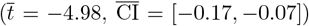. Most of these ERP differences, apart from the ones between happiness and sadness, could be explained by acoustic intensity. Furthermore, happy vocalisations evoked less positive-going ERPs than angry, neutral, and pleasant vocalisations surrounding 180 ms 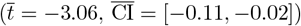, which could all be explained by acoustic intensity. In late clusters, neutral vocalisations evoked less positive-going ERPs than all others for most of the time windows after 220 ms 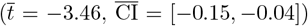, most of which could be explained by acoustic pitch and some could be explained by acoustic intensity. There were also ERP differences between anger and pleasure during 210 to 240 ms 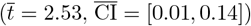, which could be explained by acoustic pitch; between happiness and pleasure during 220 and 270 ms 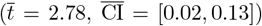, which could be partially explained by acoustic intensity. There were three clusters of ERP differences that could not be explained by intensity or pitch: angry vocalisation evoked more positive-going ERPs than happy ones during 300 to 330 ms 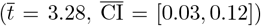 and than sad ones during 440 to 480 ms 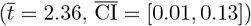; happy vocalisation evoked less positive-going ERPs than sad ones during 290 to 320 ms 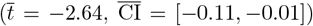. It is possible that these clusters are explainable by emotion (a point we develop in Discussion).

## 4 Discussion

In this study, we analysed 26 participants’ EEG responses to vocalisations acted in angry, happy, neutral, sad, and pleasant emotions. We analysed the effect of both vocalisations’ emotion and characteristic acoustic features (i.e., intensity, pitch, F1 and F2) on the participants’ early ERP responses. Using statistical tests done in 10 ms windows, we observed ERP differences between emotion pairs between 120 and 500 ms following stimulus onset; acoustic features, especially intensity and pitch, explained the majority of these differences. Here, we compare our findings with the literature, we consider participants’ top-down attention to emotion, and we discuss limitations of this study.

### 4.1 Important time windows and effects: agreements and disagreements with existing studies

Our analysis of EEG recordings revealed two important time windows wherein emotion appeared to affect ERP amplitudes: an early window (from 120 to 250 ms) and a late window (from 220 to 500 ms). The emotion effects we observed were mostly consistent with existing studies that also used the MAV dataset.

The first window encompassed typical P200 components. During this period, sad vocalisations evoked less positive-going ERP amplitudes as compared with the amplitudes of other emotional, and neutral, vocalisations. This response is consistent with a report by Pell et al. (2015), a study which also employed the MAV dataset. Pell et al. (2015) presented angry, happy, and sad vocalisations in a pseudo-random order to 24 healthy participants tasked with detecting emotion in facial expression pictures primed with unattended emotional vocalisations. ERPs were recorded following vocalisation onsets. The “sad” ERPs measured by Pell et al. (2015) showed less positive and later P200 components as compared to that of angry and happy vocalisations. Those authors attributed this result to the lower arousal of sad vocalisations (Pell et al., 2015). However, in our study, arousal fails to explain the less positive-going ERPs evoked by sad than neutral vocalisations because, in the MAV dataset, neutral vocalisations are less emotionally aroused than sad vocalisations (Belin et al., 2008). Rather than having to appeal to arousal, we found that most of these ERP differences could be explained by the acoustic intensity of the vocalisations. This result is consistent with previous audiological studies which reported an increase of auditory evoked P200 (P2) amplitude or N100/P200 (N1/P2) activity following simple, sinusoidal tones at higher intensity (Carrillo-de-la-Peña, 1999; Hegerl et al., 1994; Paiva et al., 2016; Sugg & Polich, 1995). In the broader literature for emotional utterance processing, changes in the P200 component amplitude have been attributed to the allocation of perceptual resources to emotionally “salient” features (Paulmann, 2023; Paulmann & Kotz, 2008; Schirmer & Kotz, 2006). Our findings neither support nor contradict this view – a plausible explanation for our finding is that, the “salient” feature is primarily the acoustic intensity, leading P200 difference to reflect the allocation of perceptual resources to intensity processing.

The second window encompassed the later stages of P200 and P300 component. Within this window, neutral vocalisations evoked less positive-going ERPs than that of emotional vocalisations. This response is consistent with a report by Pinheiro et al. (2017), describing a study which also employed the MAV dataset. Pinheiro et al. (2017) presented angry, happy, and neutral vocalisations to 19 healthy participants tasked with counting target stimuli in blocks of 250 vocalisations. The “neutral” ERPs measured by Pinheiro et al. (2017) showed less positive-going ERP amplitude (P3a) between 250 and 350 ms as compared with angry and happy vocalisations. Those authors attributed this result to arousal (Pinheiro et al., 2017). Here, we have shown that acoustic pitch and intensity could explain that effect.

### 4.2 Acoustic features provide an alternative explanation

Our results suggest an alternative explanation: that early acoustic features affect the P200 and P300 components, not the emotional content of vocalisations. Although our analysis focused on vocalisations, the early acoustic features may also account for ERP differences for other utterance types or between utterance types, as they all share the same set of acoustic features (e.g. intensity, pitch) that drives the ERPs. With this in mind we re-visited the literature describing EEG studies of emotional speech/vocalisation perception. But re-examination of this literature was not straightforward because most studies report only derived acoustic features averaged across entire stimulus recordings, thus obscuring any association between acoustic features and the P200 and P300 components.

An exception is the above-mentioned study by Pell et al. (2015). Those authors presented both emotional pseudo-sentences (e.g., “He placktered the tozz”) and emotional vocalisations drawn from the MAV dataset (anger, happiness, and sadness), examining averaged ERP amplitudes separately for the pseudo-sentences and vocalisations, and for emotion. Critically, they reported the intensity and pitch of stimuli just prior (between 0 and 160 ms relative to stimulus onset) to their analysis of ERP amplitude (between 170 and 300 ms) over the fronto-central scalp region.

Pell et al. (2015) observed a greater P200 amplitude in response to sad and angry vocalisations than that of pseudo-sentences, and attributed this effect to a “rapid deployment of attentional resources” for emotional salience detection . Our findings provide an alternative explanation. Pell’s sad vocalisations had higher pitch (between 0 to 160 ms) then sad pseudo-sentences (289 vs 230 Hz), and Pell’s angry vocalisations, compared to pseudo-sentences, had both a higher pitch (317 vs 214 Hz) and a greater intensity (74 vs 69 dB) (Perrin et al., 1989). These acoustic differences could thus explain the effects on P200 amplitude. Similarly, Pell et al. (2015) observed a greater P200 amplitude following happy pseudo-sentences as compared to angry pseudo-sentences, but not for happy and angry vocalisations. Again, our findings provide an explanation: Pell’s happy pseudo-sentences had both a greater intensity (75 dB) and higher pitch (250 Hz) than angry pseudo-sentences (intensity, 69 dB; pitch, 214 Hz), but this was not the case for Pell’s happy and angry vocalisations (Pell et al., 2015). These acoustic differences could thus explain these effects on P200 amplitude.

### 4.3 Participants’ top-down attention to emotion

In our study, participants were instructed to recognise the emotion category and then reproduce the /ah/ vowel in the recognised emotion. They thus applied top-down attention to emotion while listening to emotional vocalisations. From the second block onwards, as participants became increasingly familiar with the stimuli (the same set was used across blocks), fewer attentional resources may have been directed to emotion but more toward the acoustic aspect.

The effect of top-down attention on auditory-evoked ERP amplitudes is well documented and broadly accepted (Hansen & Hillyard, 1983; Hillyard et al., 1973; Hink et al., 1978; Tervaniemi et al., 2009). Attentional effects are also observed during the perception of human utterances. Power et al. (2012) demonstrated that top-down selective attention enhances the cortical response that continuously tracks the acoustic features of temporally unfolding utterances. In this regard, increased attention to acoustic features may have enhanced the observed acoustic-ERP correlations. However, this enhancement in Power et al. (2012) was observed in a dichotic setting, where participants attended one of the two audio streams presented to different ears. In our study, participants attended a single audio stream, leaving open the question of whether top-down attention to specific acoustic features within the stimulus would similarly enhance the cortical response tracking the stimulus.

The potential reduction in top-down attention to emotion may have reduced the apparent emotion effects on early ERPs (i.e., P200, P300) in favour of an acoustic-driven explanation. Although participants’ top-down attention to emotion has been proposed to affect the neural mechanisms involved in emotional utterance perception (Paulmann, 2023; Schirmer & Kotz, 2006), limited ERP evidence is available. To our knowledge, only one study by Wambacq et al. (2004) reported a marginally more positive-going P300 during attentive than inattentive processing of negative prosody in spoken words. This difference, however, did not reach statistical significance (p=0.06). In contrast, the literature has repeatedly reported emotion effects on early ERP components (e.g., P200, P300, N400, LPP) regardless of whether the participant was attentive to the emotion aspect of stimuli or not (Gädeke et al., 2013; Paulmann et al., 2013; Pell et al., 2015; Pinheiro et al., 2015; Steber et al., 2020).

### 4.4 Limitations and future directions

In our study, we limited the analysis to four characteristic acoustic features (i.e. intensity, pitch, first formant and second formant), individually assessing their correlation with the ERPs. This selection allows a comprehensive understanding of the ERP effects induced by key acoustic features, but raises several limitations. There exist additional acoustic features that were not analysed in our study. Moreover, we computed the acoustic features within 10 ms windows and did not compute the temporal variability of the features (e.g., the peak-to-peak amplitude of pitch in 100 ms windows). Furthermore, emotions in utterances are usually distinguished by the integration of multiple acoustic features (Schirmer & Kotz, 2006), while the complex interactions between acoustic features were not considered in our analysis. In future studies, carefully designed analysis frameworks with more advanced statistical or machine learning tools are required to address these complex, multifactorial acoustic features.

Our finding that acoustic features account for early ERP differences does not confirm or refute an abstraction of emotion representation in the brain during this early perception period. In future studies, more advanced statistical analysis methods such as representational similarity analysis (RSA) (Kriegeskorte et al., 2008) might be used to identify emotion abstraction after accounting for the effects of temporally dynamic acoustic features. In a recent study, Lavan et al. (2024) used RSA to examine the time course for the abstraction of person characteristics (e.g., gender, attractiveness, educatedness, etc.) during human voice perception, after accounting for acoustic differences in stimuli. The authors, however, again used acoustic features averaged across entire stimulus recordings and lacked considerations of the temporal dynamics of the acoustic features. In future research, to extend this method with consideration for the temporal dynamics, a much more complicated and richer computation for acoustic features in the RSA model is required.

In our results, we also observed emotion effects not explained by the four characteristic acoustic features (i.e. intensity, pitch, first formant and second formant). The acoustic features might have failed to explain these effects due to a lack of statistical power in verifying the associations between these acoustic features and the ERP amplitudes, or the acoustic differences between pairs of emotion categories. The failure might also be due to interactions between these acoustic features. However, it is possible that some of these emotion effects may in fact reflect the neural processing of emotion per se. It is noteworthy that, across actors, happy vocalisations shared similar intensity with sad vocalisations, which was lower than that of other emotions. Despite their low acoustic intensity, happy vocalisations evoked a more positive-going ERPs than that of sad vocalisations between 120 and 250 ms with a relatively high robustness 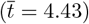. This ERP difference could not be explained by intensity alone, nor could it be satisfyingly explained by pitch, F1, or F2. This result indicated a potential difference in brain’s processing of happy vocalisations as compared to others.

In this study, we focused on addressing mixed results in the literature on emotion effects for early ERPs. We thus restricted the ROI to fronto-central channels, where the N100, P200 and P300 components peaked. In this study, ERP amplitudes over other relevant cortex regions were not considered. In particular, the auditory cortex, the superior temporal gyrus (STG), and the anterior superior temporal sulcus (STS) under the temporal regions were theorised to be involved in processing and integrating acoustic features during the perception of emotional utterances (Schirmer & Kotz, 2006). Future studies targeting these regions might help reveal more details of the underlying brain mechanisms in processing acoustic features in emotional utterances.

## 5 Conclusion

In summary, we recorded ERP responses of participants during their perceptions of emotional vocalisations. We analysed instantaneous intensity, pitch, F1, and F2 acoustic features in conjunction with ERP amplitudes, and emotion categories. ERP differences between emotion categories emerged across multiple time windows between 120 and 500 ms post-stimulus onset, over the fronto-central region of the scalp, while most of these emotion effects could be explained by acoustic differences 100 ms prior to the corresponding window. Further analysis and comparisons with existing literature suggested impacts of the intensity and pitch on the amplitudes of early ERP components (i.e., P200 and P300). These findings serve as a reminder to researchers interested in the neural correlates of emotion to account for the effects of stimulus features before making inferences about emotional processing. Our data contribute to a deeper understanding of the nuanced role played by key acoustic features in the early stages of the perception of emotional utterances, and could help explain the differences in emotion effects reported in the literature.

## CRediT Author Statement

**Yichen Tang**: Conceptualization, Data curation, Formal analysis, Investigation, Methodology, Resources, Software, Validation, Visualization, Writing – original draft, Writing – review & editing. **Paul M. Corballis**: Conceptualization, Methodology, Supervision, Writing – review & editing. **Luke E. Hallum**: Conceptualization, Methodology, Funding acquisition, Project administration, Resources, Supervision, Writing – review & editing.

## Data and code availability statement

The experimental data generated during the current study are not publicly available due to privacy concerns. However, the processed data and related code that support the findings of this study can be made available on request. Requests for access will be reviewed on a case-by-case basis to ensure they comply with ethical standards.

## Ethics approval statement

The experimental protocols have been approved by the University of Auckland Human Participants Ethics Committee (ref. UAHPEC24522).

