## Appendices for "Acoustic features of emotional vocalisations account for early modulations of event-related brain potentials"

### Appendix A

### Extracted and Corrected Pitch Contours

Figures A-1 to A-5 show the original waveform, the extracted pitch contour using Praat with manual corrections, and the spectrogram of five sample vocalisations - anger, happiness, neutral, pleasure, and sadness - performed by Actor 6 from the Montreal Affective Voices (MAV) dataset. Figures A-6 and A-7 show the extracted and corrected pitch contours for all vocalisations performed by the remaining actors. The vocalisations whose pitch contours were manually corrected are: 6_happiness, 45_sadness, 46_happiness, 46_neutral, 53_neutral, 53_sadness, 55_happiness, 58_pleasure, 58_sadness, 59_happiness, 59_pleasure, 60_happiness, 60_neutral, 60_pleasure, and 61_sadness.

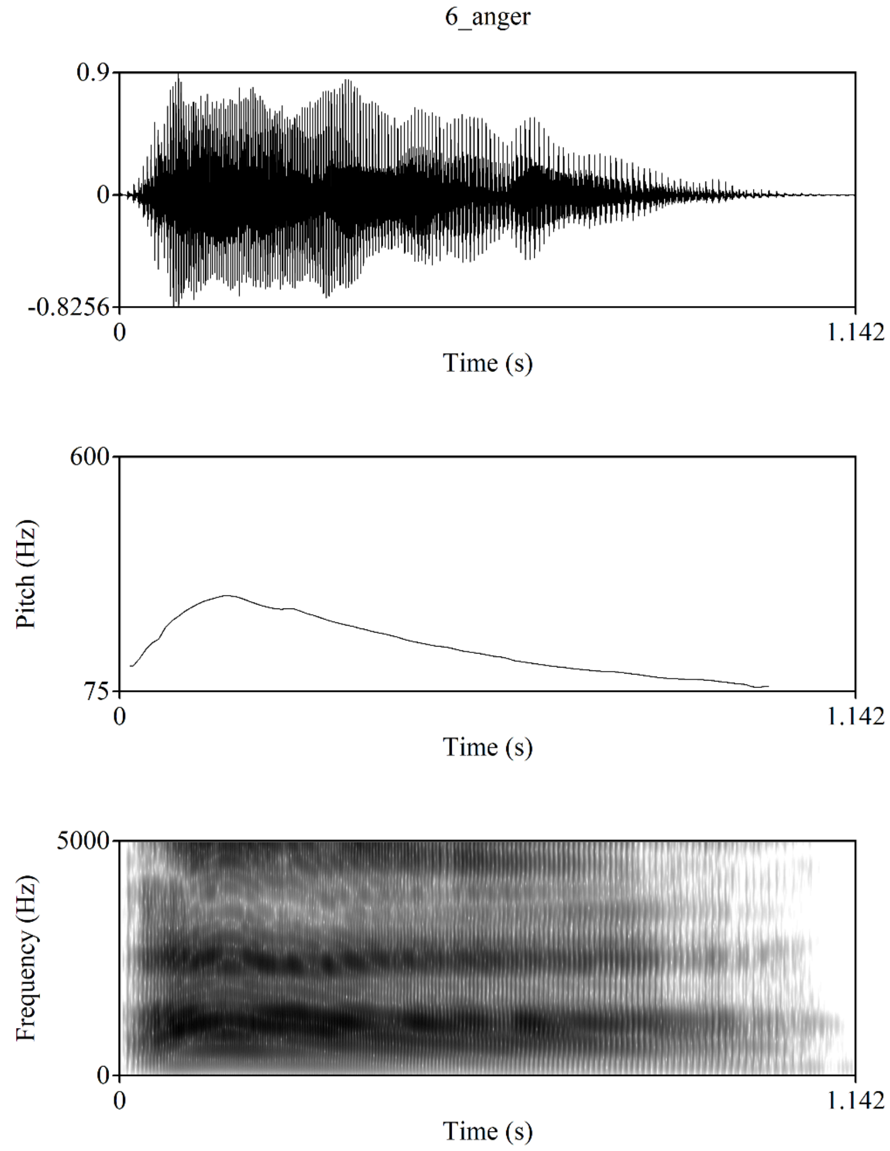


**Figure A-1** The waveform, pitch contour, and spectrogram of the anger vocalisation produced by Actor 6 from the Montreal Affective Voices (MAV) dataset.

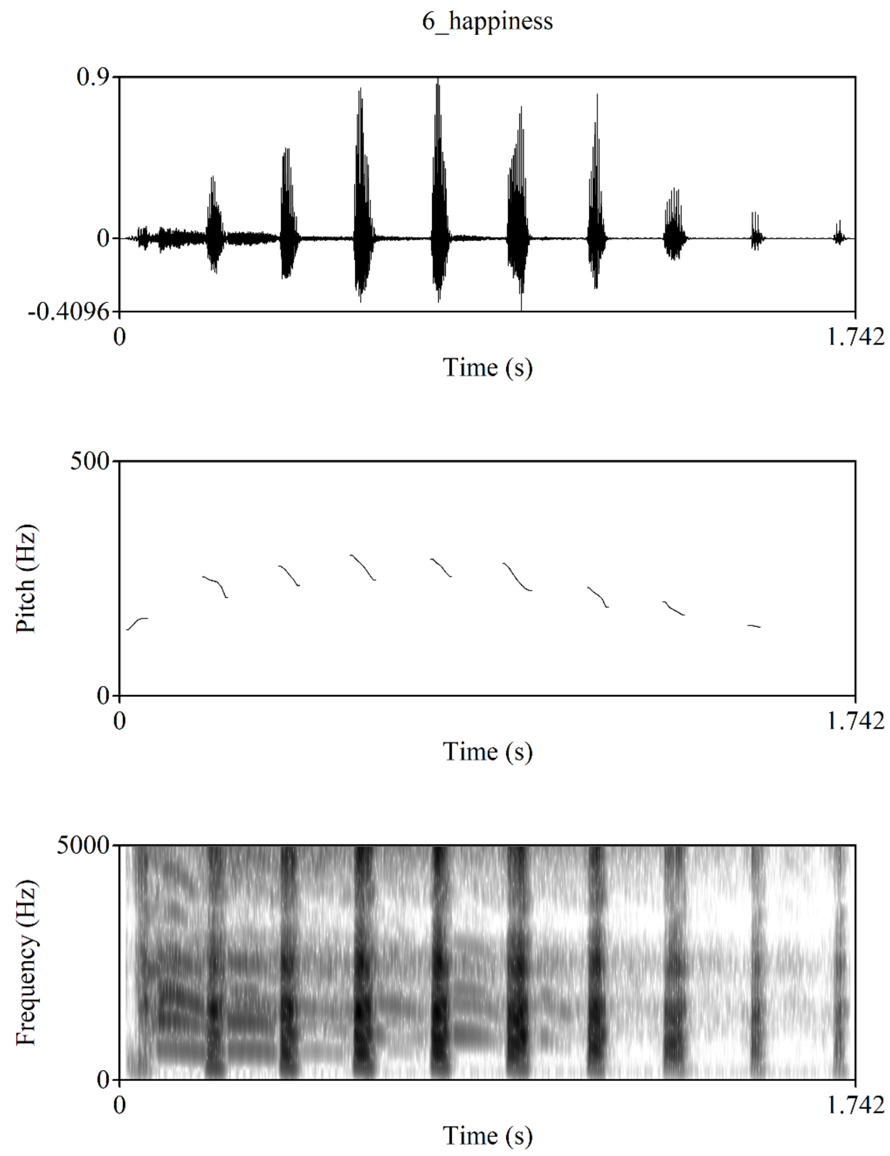


**Figure A-2** The waveform, pitch contour, and spectrogram of the happiness vocalisation produced by Actor 6 from the Montreal Affective Voices (MAV) dataset.

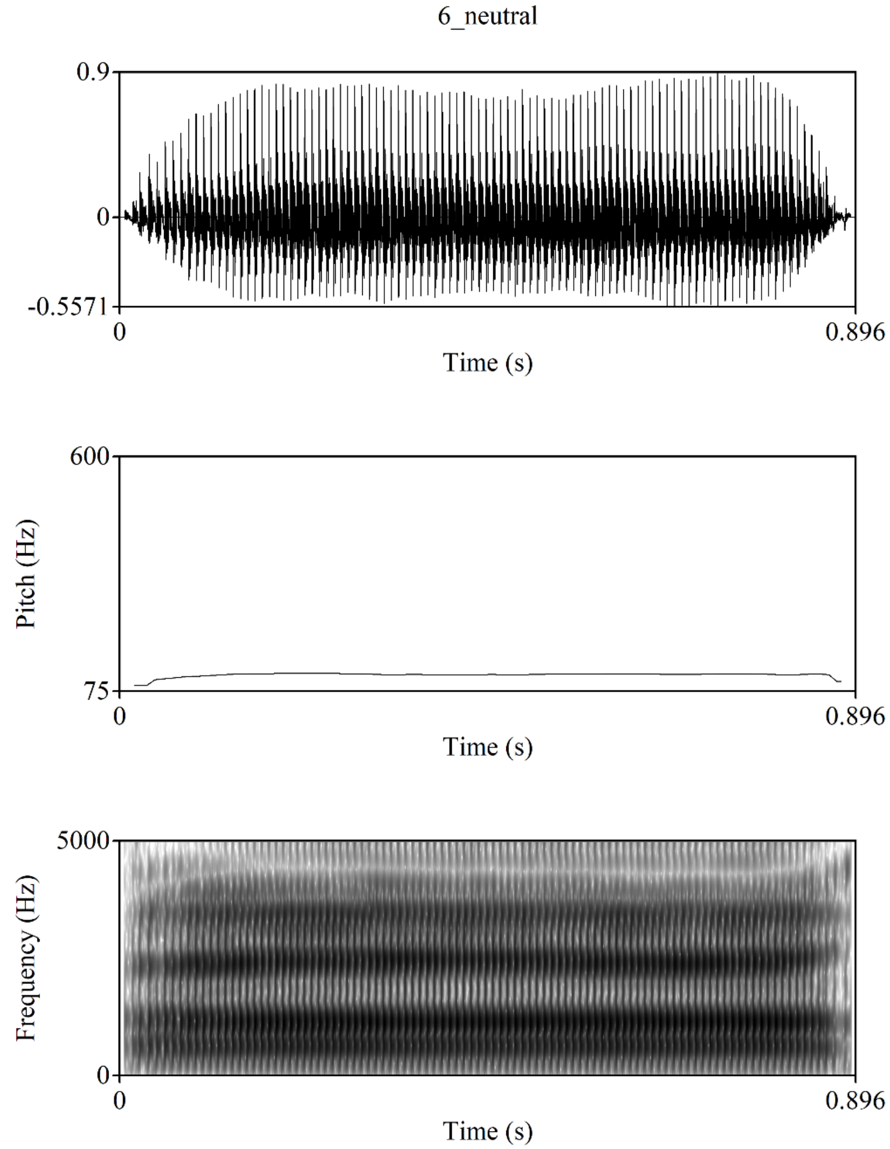


**Figure A-3** The waveform, pitch contour, and spectrogram of the neutral vocalisation produced by Actor 6 from the Montreal Affective Voices (MAV) dataset.

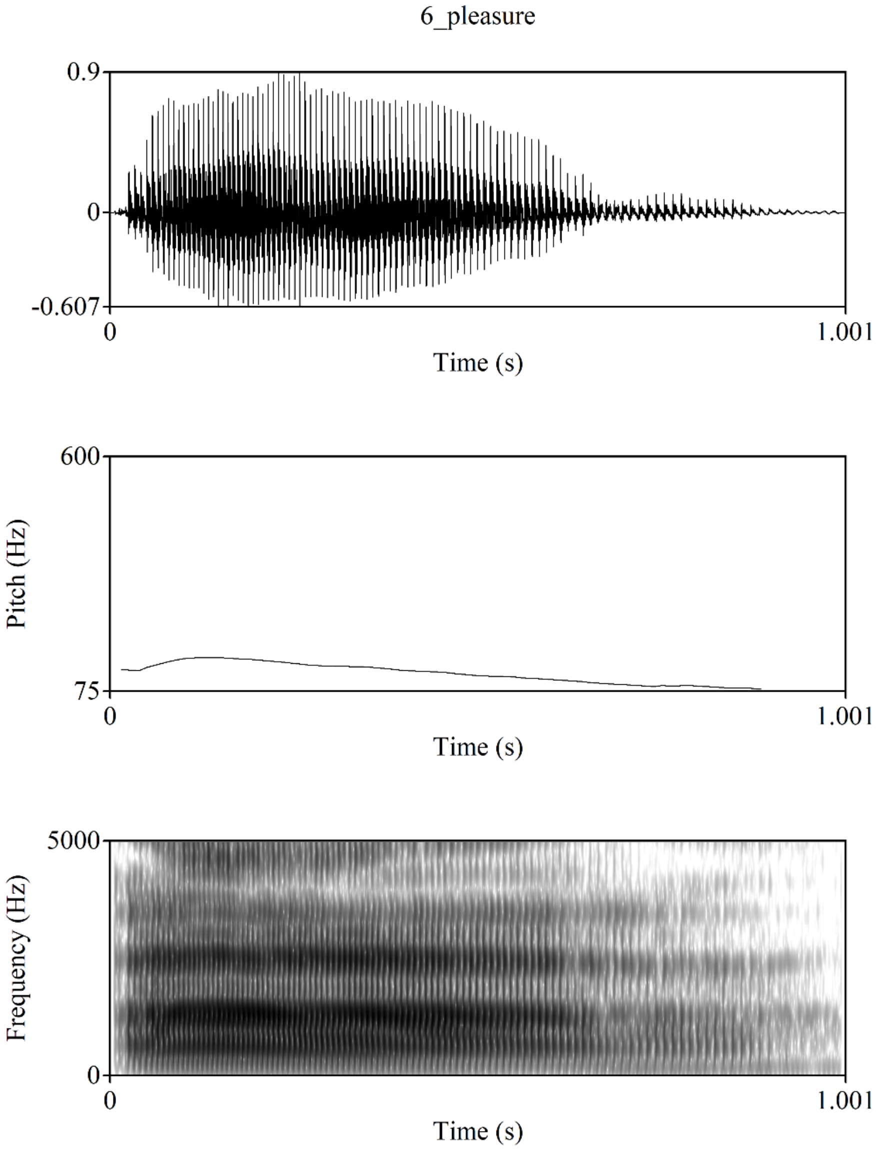


**Figure A-4** The waveform, pitch contour, and spectrogram of the pleasure vocalisation produced by Actor 6 from the Montreal Affective Voices (MAV) dataset.

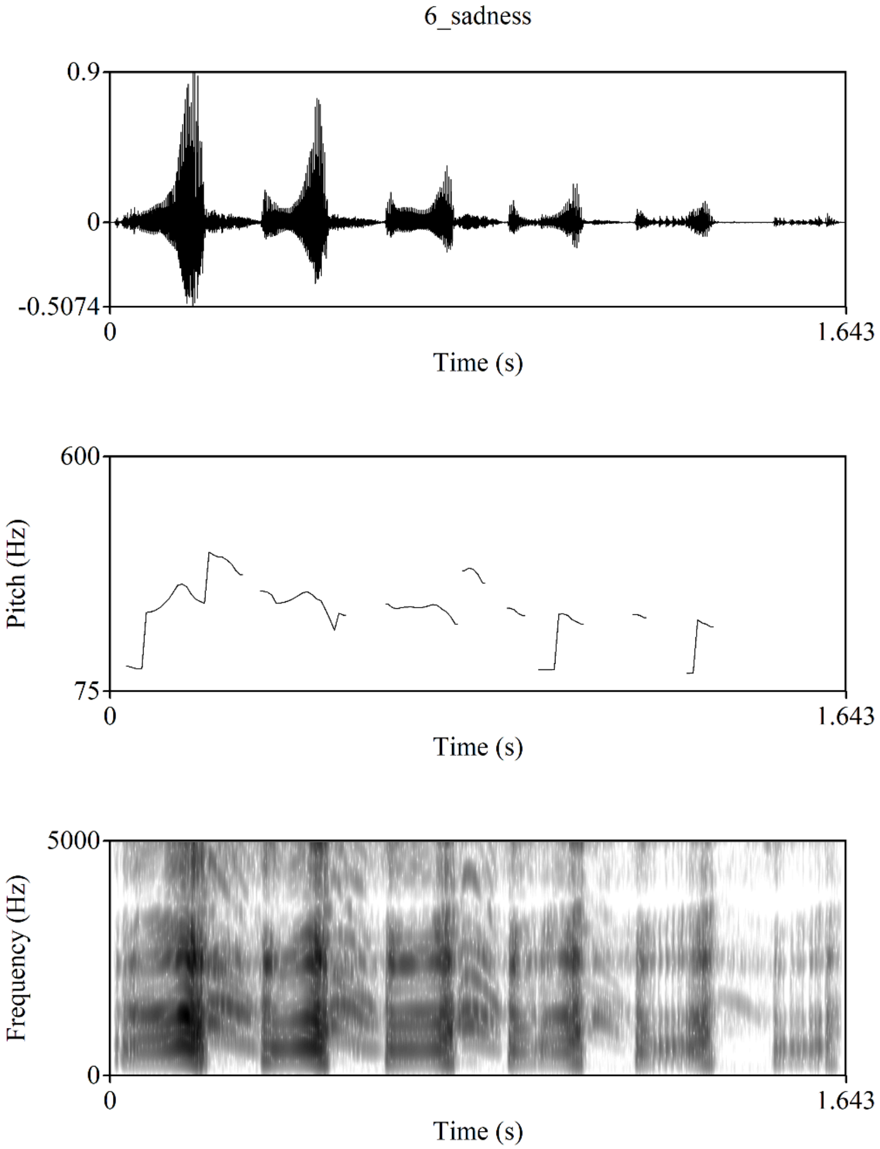


**Figure A-5** The waveform, pitch contour, and spectrogram of the sadness vocalisation produced by Actor 6 from the Montreal Affective Voices (MAV) dataset.

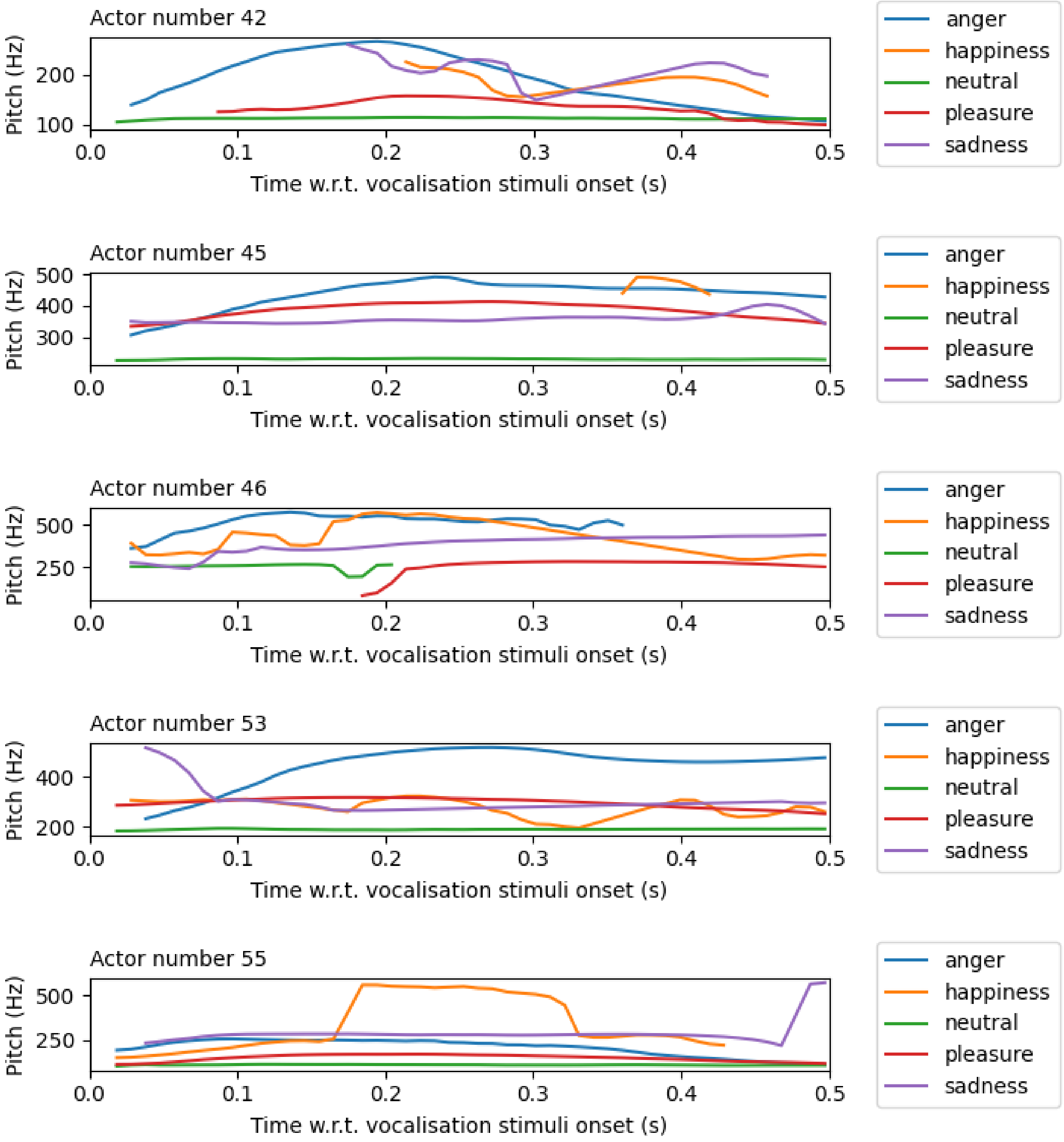


**Figure A-6** The pitch contours of vocalisations performed by Actors 42, 45, 46, 53 and 55 from the Montreal Affective Voices (MAV) dataset.

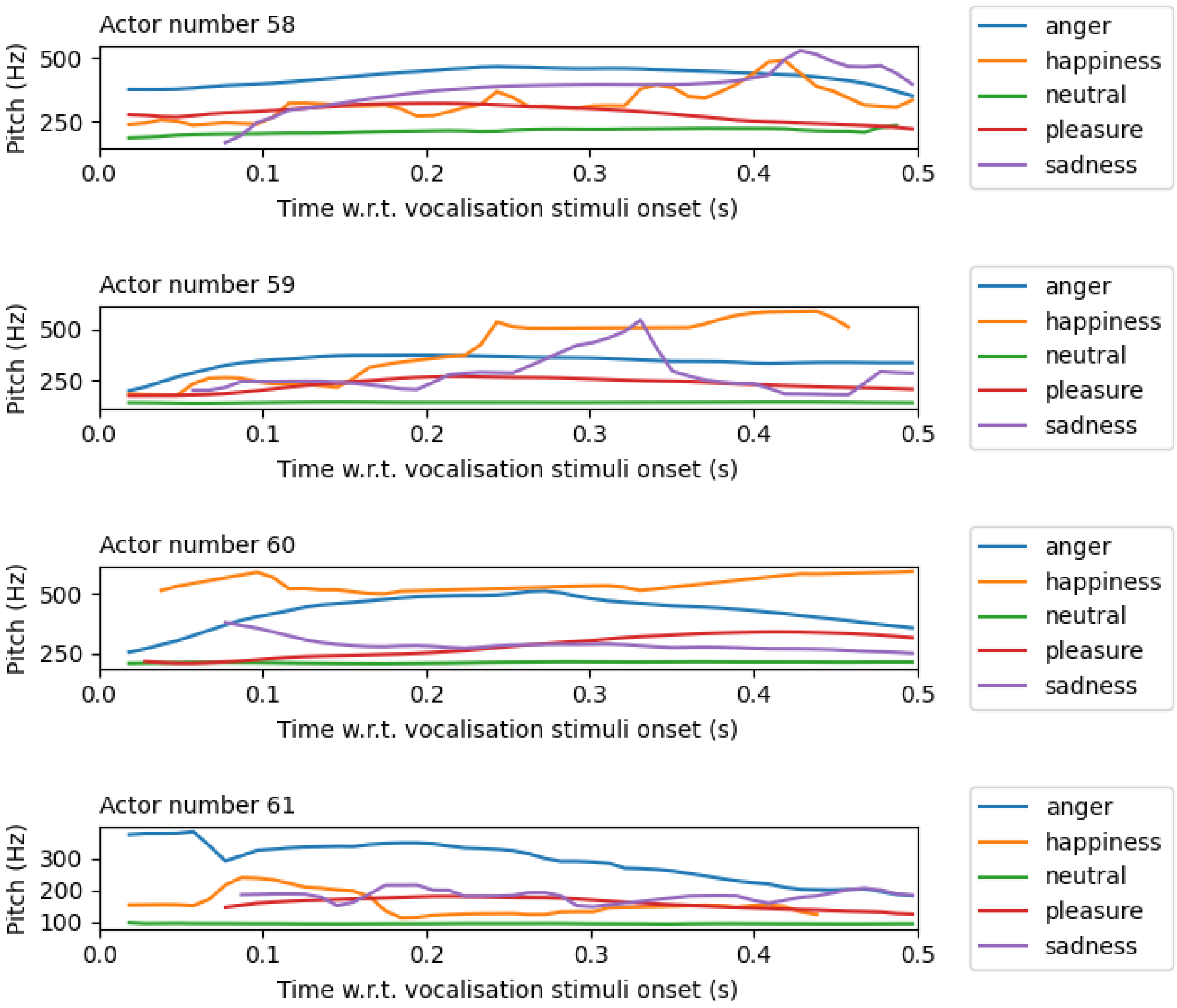


**Figure A-7** The pitch contours of vocalisations performed by Actors 58, 59, 60 and 61 from the Montreal Affective Voices (MAV) dataset.

### Appendix B

#### Differences in instantaneous harmonic-to-noise ratio (HNR) between emotion category pairs

We computed the instantaneous harmonic-to-noise ratio (HNR) for all 50 vocalisations using Praat, with a step size of 10 ms. Figure B-1 shows the instantaneous HNR for five sample vocalisations performed by Actor 55 from the MAV dataset. Differences in HNR between emotion pairs were evaluated using two-sided, two-sample Student's $t$-tests, as described in Section 2.2.3. Statistically significant differences, along with the corresponding t-statistics, are shown in Figure B-2. A 100 ms lag was added to the time axis, as described in Section 2.2.3, to align it with Figure 7.

HNR differences were observed in only four emotion pairs across 37 time windows. Although the mean t-statistics were high ($|\bar{t}| = 7.90$, $\overline{|CI_{95}|}\ for\ mean\ HNR\ difference = [47.94, 130.45]$), differences in most emotion pairs were confined to short time intervals: anger vs. neutral, 0.12-0.31 s, $\bar{t} = 3.03$, $\overline{CI} = [3.30, 18.66]$; happiness vs. pleasure, 0.28-0.33 s, $\bar{t} =- 12.46$, $\overline{CI} = [- 263.94, - 96.14]$; anger vs. happiness, 0.28-0.33 s, $\bar{t} = 13.31$, $\overline{CI} = [98.55, 266.19]$; and happiness vs. neutral, 0.29-0.33 s, $\bar{t} =- 15.40$, $\overline{CI} = [- 254.50, - 107.89,]$.

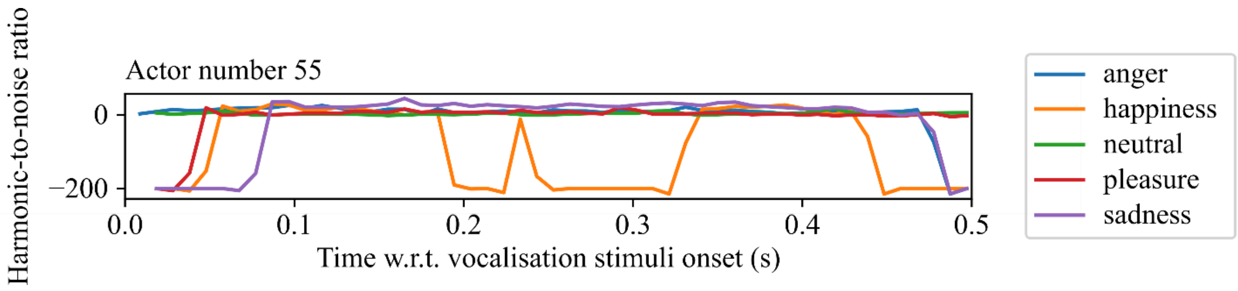


**Figure B-1** The instantaneous harmonic-to-noise ratio (HNR) measured for five emotional vocalisations produced by Actor 55 from the Montreal Affective Voices (MAV) dataset.

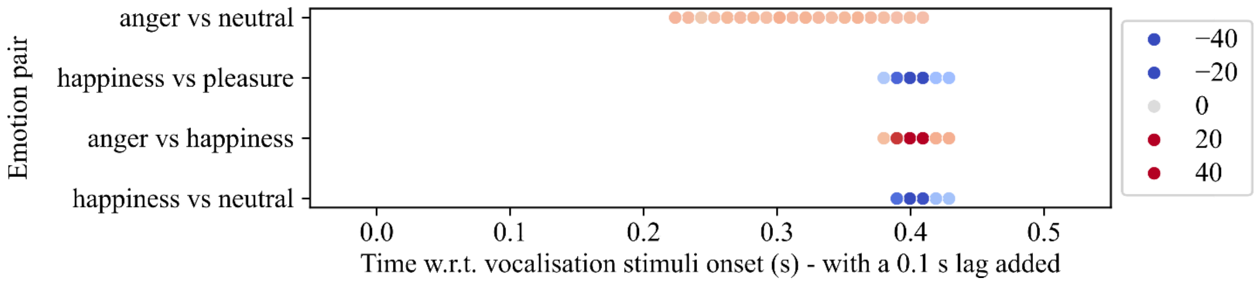


**Figure B-2** The points in time where there were statistically significant differences in instantaneous HNR between emotion category pairs. The red/blue colour map represents the value of the t statistic (see Methods). Here, we adjusted the x-axis to account for the 100 ms lag used. For example, the HNR of angry vocalisations was statistically significantly greater than that of happy vocalisations during 280 to 330 ms ($\bar{t} = 13.31$, $\overline{CI_{95}}\ for\ mean\ HNR\ difference\ = [98.55, 266.19]$, plotted as 380 to 430 ms on the x-axis). Emotion pairs with no statistically significant differences in acoustic features are omitted.
